# Lysogen formation governs colonies while lytic infection is more prevalent in single cells of the bloom-forming cyanobacterium, *Microcystis*

**DOI:** 10.1101/2025.02.12.637950

**Authors:** X. Huang, EE Chase, BN Zepernick, RM Martin, LE Krausfeldt, HL Pound, H. Wu, Z. Zheng, SW Wilhelm

## Abstract

While the bloom-forming cyanobacterium *Microcystis* can exist as free-living single cells or within dense mucilaginous colonies, the drivers and consequences of colony formation remain unclear. Here, we integrated metatranscriptomic datasets from two *Microcystis* bloom events in Lake Taihu, China, to analyze and validate the functional differences between colonial and single-cell *Microcystis*. Our results confirmed colony expression profiles were disproportionately enriched in *Microcystis* transcripts (and functions) compared to other prokaryotic taxa. Concomitantly, viral infection strategies diverged by *Microcystis* community morphology: colony-associated cells expressed lysogeny-associated genes, while single cells exhibited increased signatures of lytic infection. These data are consistent with the hypothesis that *Microcystis* colonies foster conditions favorable to lysogen formation—likely due to local high cell densities and the resulting advantage of superinfection immunity—whereas solitary cells experience stronger lytic pressure. On a broader scale, our findings refine the understanding of bloom dynamics by identifying how community morphological states coincide with distinct host–virus interactions. Cumulatively, this work underscores the importance of colony formation in shaping *Microcystis* ecology and highlights the need for mechanistic studies that disentangle the interplay between phage infection modes, colony formation, and microbial community structure.

## Introduction

Cyanobacteria are among the most ecologically significant microorganisms in fresh waters, driving global biogeochemical cycles that include carbon fixation and nutrient dynamics. Yet, concomitantly with these benefits come considerable environmental challenges when these organisms form harmful algal blooms^1^. *Microcystis* has emerged globally as a key player^2^ due to its adaptability to environmental conditions^3^. While colony formation is a hallmark of *Microcystis* in natural systems, this capability is often lost under laboratory conditions^4^, underscoring the potential importance of environmental pressures in driving this phenomenon.

The formation of *Microcystis* colonies has been suggested to confer numerous ecological advantages, including resistance to chemical stress^5^, high-light conditions^6^, and grazing pressure^7^, while facilitating buoyancy and nutrient acquisition^6^. Central to many of these processes is the interaction between *Microcystis* and associated heterotrophic bacteria (*i*.*e*., microbiome or phycosphere community), which can be embedded within the extracellular polymeric substance (EPS) matrix surrounding colonies^8^. This phycosphere community, comprising diverse taxa, is hypothesized to play pivotal roles in nutrient cycling, organic matter decomposition, and resource exchange, creating a microenvironment that supports *Microcystis* proliferation and dominance in eutrophic waters^9^.

Bacteria and phages represent the most abundant and genetically diverse entities on Earth, with phages often outnumbering their bacterial hosts by an order of magnitude^10^. Many phages employ a dual strategy of infection: entering lytic or lysogenic cycles^10^. In the lysogenic state, phages integrate into the bacterial genome as prophages, establishing a symbiotic relationship that may impose a fitness cost by disrupting host gene expression but also potentially conferring adaptive advantages, such as regulating host gene expression^11^, introducing or altering functions^10^, facilitating bacterial DNA transfer^12^, and shaping bacterial communities^13–15^. These phage-mediated interactions are thought to facilitate functional and evolutionary shifts in bacterial hosts, particularly under environmental stress^16,17^.

In this study, we used 20- and 0.45-μm-pore-size filters to enrich for colonial or single-celled *Microcystis* in bloom samples from Lake Taihu, China, and analyzed metatranscriptomic data to investigate the interplay between colony formation, the associated microbiome and viral infection. Additionally, we used data from another year (collected but not processed) to validate the experimental results. Previously we had hypothesized that lysogen formation or growth at high cell densities would be more prevalent when cyanobacterial blooms were at high densities^18^: we thought resistance to superinfection commonly seen in other lysogens might protect these massive blooms from collective lysis due to the high virus-host contact rates conditions would create^19^. By profiling the expression of lysogeny- and lytic-cycle-associated genes in a natural system, we demonstrated a striking correlation between colony formation and lysogenic activity. Our findings highlight the complexity of *Microcystis* ecological strategies and underscore the need for further research, including controlled experiments in laboratory settings, to disentangle the mechanistic links between lysogeny, colony formation, and microbial interactions.

## Methods

### Sample collection

Surface water samples of a *Microcystis* bloom were collected from four sites (Supplementary Table 1) in Zhushan Bay, Lake Taihu (*Taihu* in Mandarin) on August 26, 2023 at midday The separation of single cells and colonies of *Microcystis* was completed by a vacuum filter device (pressure range: 0 to -1 MPa) equipped with 20 μm filter membranes. Subsequently filtrates were pass through 0.45 µm filters to collect single cells. At each of four locations, sampling was performed in triplicate, resulting in a total of 12 colonial samples and 12 single-cell samples. Immediately after separation, the filter membranes were stored at -80° C until further processing.

In parallel with the above samples, we accessed previously collected and sequenced data for samples taken in 2018. Six colonial and six single-cell samples were collected from the boat dock of the *Taihu Laboratory for Lake Ecosystem Research*. Samples were collected using 28 µm mesh-size Nytex™ filtration material: retentate was maintained for the “colonial” size class while material passing through the filter was recollected onto a 0.2-μm nominal pore-size polycarbonate filter mounted in a Swinnex™ holder and delivered with a sterile 60 CC syringe. Sample collections for 2018 dataset are detailed in Supplementary Information.

### RNA extraction and sequencing

RNA samples were extracted and sequenced at Shanghai Majorbio Bio-pharm Biotechnology Co., Ltd. (Shanghai, China). Total RNA was extracted from the tissue using Soil RNA Extraction Kit (Majorbio, China). Total RNA was processed using the Illumina® Stranded mRNA Prep, Ligation kit (Illumina, San Diego, CA, USA), with rRNA depletion performed using the RiboCop rRNA Depletion Kit for mixed bacterial samples (Lexogen, USA). Libraries were prepared following standard Illumina protocols and sequenced on an Illumina NovaSeq 6000 platform (paired-end mode). Sample collection, RNA extraction and sequencing of 2018 samples are detailed in Supplementary Information.

### Pangenome assembly

To identify the transcriptomic differences between *Microcystis* morphotypes, 16 complete, closed *Microcystis* genomes (Supplementary Table 2) were downloaded from the National Center for Biotechnology Information (NCBI) to establish a *Microcystis* pangenome^20^. All individual genomes were merged into a single file and redundant coding sequences were subsequently removed *via* CD-HIT (nucleotide identity of 0.95) (v.4.8.1)^21^. Functional annotation of the *Microcystis* pangenome was derived from the individual genomes, which were automatically annotated using NCBI Prokaryotic Genome Annotation Pipeline (PGAP)^22^. Additional annotations were supplemented using EggNOG-mapper (v.2.1.12)^23^ using a specified e-value of 1e^-10^.

### Metatranscriptomic analyses

The sequence analyses of both 2023 and 2018 libraries used the following pipeline: bioinformatic trimming of raw reads was performed using fastp (v.0.23.2)^24^. Subsequently, reads were interleaved together using reformat.sh script available in the BBTools suite(v.38.18)^25^. Residual rRNA and contaminants were removed using the JGI reference database and bbmap.sh(v.38.18)^25^ (minid=0.93). The quality of reads was checked using FastQC (v.0.12.1) before and after trimming. Trimmed and filtered libraries (n=24) were concatenated and assembled via MEGAHIT (v.1.2.9)^26^, with quality of coassembly confirmed via QUAST QC (v.5.0.2)^27^. Read mappings were performed using bbmap.sh (v.38.18)^25^ (minid= 0.93) to align mRNA reads to the coassembly, *Microcystis* pangenome and Ma-LMM01 genome, respectively. The summary of sequence information of 2023 libraries was listed in Supplementary Table 3. Read counts were tabulated by featureCounts (v.2.0.6)^28^.

Gene predictions were performed using MetaGeneMark (v.3.38)^29^ with the metagenome-style model. Taxonomic annotation of predicted genes was performed using Kraken 2 (v.2.1.3)^30^ with the RefSeq complete genomes dataset (including archaea, bacteria, viral, fungi, and plant genomes). Functional annotations of predicted genes were supplemented using EggNOG-mapper (v.2.1.12)^23^ with a specified e-value of 1e-10. Raw sequencing data for all 24 transcriptomic libraries are available at the NCBI Sequence Read Archive (SRA) under the accession number PRJNA1206705.

### Statistical analyses

Mapped reads were normalized to transcripts per million (TPM), representing relative transcript abundance, to account for differences in library size and gene length. The terms “overrepresented” and “underrepresented” have been used to describe changes in the proportional representation of transcripts within the total transcriptome pool across conditions. All analyses were conducted in R^31^ (v.4.4.2). Normality was assessed, and paired t-tests were applied for normally distributed data, while Wilcoxon signed-rank tests were used for non-normal data. Clustering of the normalized libraries was visualized *via* nonmetric multidimensional scaling (NMDS) using Bray–Curtis dissimilarity, implemented in the vegan (v.2.6-8) package. KEGG pathway enrichment analysis was performed using the clusterProfiler^32^ (v.4.14.4) package. Heatmaps were generated with the pheatmap (v.1.0.12) package, and all additional figures were created using the ggplot2^33^ (v.3.5.1) package. Differential expression analysis was conducted using DESeq2^34^ (v.1.28.1), with genes showing a baseMean < 10 excluded as noise. Genes with a log_2_ fold change > 1 (log_2_|FC| >1) and an adjusted *p* value of < 0.05 were considered differentially expressed.

## Results

### Colonies were less taxonomically diverse than single-cell samples

We observed significant differences in the taxonomic composition between colonial (Fig. 1a) and single-cell (Fig. 1b) samples. *Cyanobacteriota* dominated both colonial and single-cell samples, accounting for an average of 94.7% in colonial samples and 72.1% in single-cell samples, with a significantly higher proportion in the colonial samples (*p* < 0.001, paired *t*-test). Excluding *Cyanobacteriota*, the top five phyla which all demonstrated lower relative abundance in colonial samples compared with single-cell sample and included *Pseudomonadota* (*p* < 0.001, paired *t*-test), *Bacteroidota* (*p* < 0.001, paired *t*-test), *Actinomycetota* (*p* < 0.001, paired *t*-test), *Streptophyta* (*p* < 0.001, paired *t*-test) and *Bacillota* (*p* < 0.001, paired *t*-test). At the genus level, *Microcystis* was the most abundant within the phylum *Cyanobacteriota*, accounting for an average of 91.7% in colonial samples and 67.9% in single-cell samples (Supplementary Figure 1). During the 2023 sampling, *Microcystis* overwhelmingly dominated both large colonies and small particles like *Cyanobacteriota* did at phylum level, while other bacteria and eukaryotes were present at concentrations 1–2 orders of magnitude lower.

**Fig. 1.**
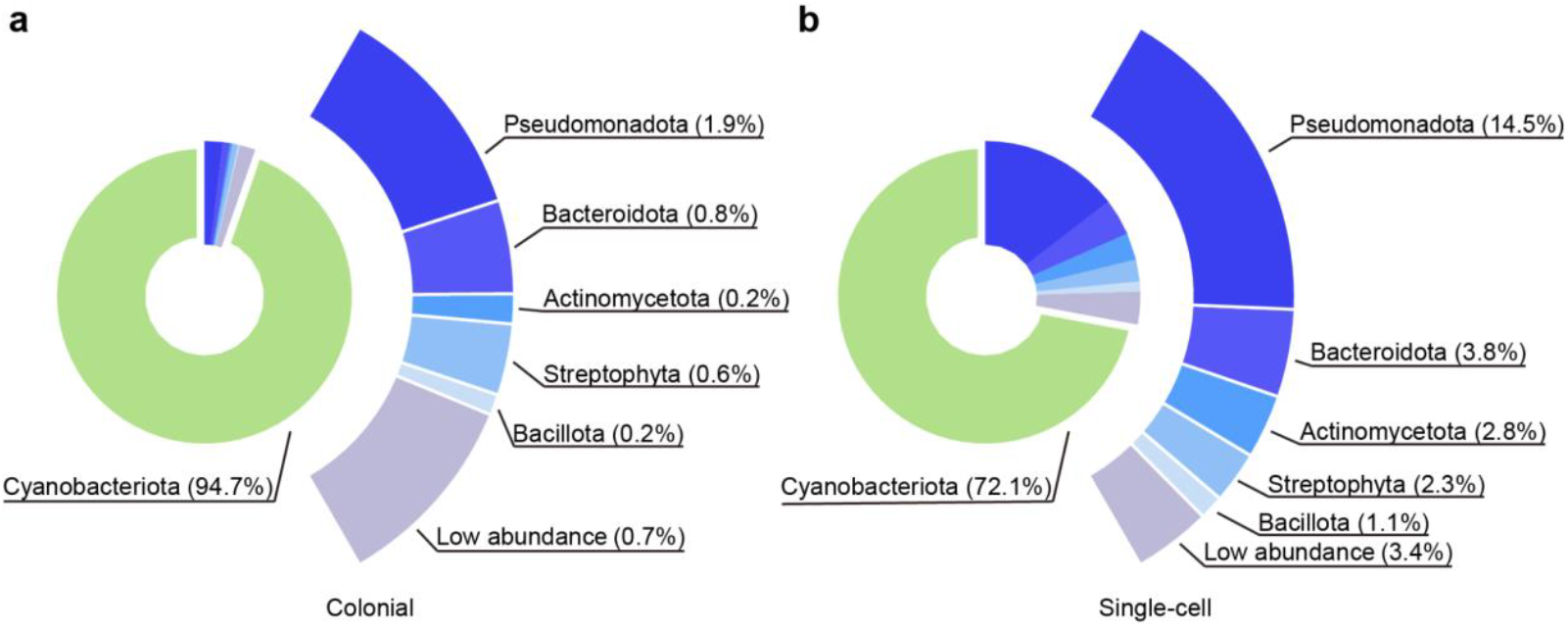
Microbiota of colonial and single-cell samples. **a, b**, Phylum-level distribution of the microbiota in colonial samples (**a**, n = 12) and single-cell samples (**b**, n = 12). *Cyanobacteriota* are represented in the inner donut chart. The outer ring highlights the relative abundances of non-*Cyanobacteriota* phyla, with corresponding percentages indicated beside the taxa.

### *Microcystis* dominated transcription of biological processes in colonies

Gene expression of both the microbiota and *Microcystis* differed in the colony samples relative to single-cell samples. The nMDS of Bray-Curtis distance based on gene expression from both the co-assembly (Fig. 2a) and *Microcystis* pangenome (Fig. 2b) revealed that colonial and single-cell samples formed two distinct clusters. Table 1 presents the results of the differential expression (DE) gene analysis among the top six phyla mentioned before. In *Cyanobacteriota*, overrepresented and underrepresented genes between colonies and single cells were comparable. However, in other bacteria and eukaryotes, 97.8%–99.9% of the relevant expression of the genes were underrepresented in colonial samples. Within *Microcystis* alone, 81.62% of the genes in *Microcystis* increased expression in colonial libraries, whereas 97.76% of the genes in other *Cyanobacteriota* decreased expression. This latter pattern was similar to that of other bacteria and eukaryotes. The KEGG pathway enrichment analysis genes with decreased expression among all microbiota, excluding *Microcystis*, were primarily enriched in biosynthesis of cofactors and amino acids, and carbon metabolism (Supplementary Figure 2). Within these same pathways, *Microcystis* showed more genes with increased expression (n = 212) than decreased expression (n = 116). (Supplementary Figure 2)

**Table 1.** Number of DE genes based on the co-assembly and annotated by KEGG.

| Taxonomy | Total number of DE genes | Number of overrepresented genes in colonies | Number of underrepresented genes in colonies |
| --- | --- | --- | --- |
| *Cyanobacteriota* | 10,509 | 4,423 | 6,086 |
| *Microcystis* | 5,404 | 4,346 | 1,058 |
| *Cyanobacteriota*<br>excluding *Microcystis* | 5,105 | 77 | 5,028 |
| *Pseudomonadota* | 54,612 | 66 | 54,546 |
| *Bacteroidota* | 17,150 | 23 | 17,127 |
| *Actinomycetota* | 11,515 | 13 | 11,502 |
| *Streptophyta* | 4,801 | 117 | 4,684 |
| *Bacillota* | 2,491 | 16 | 2,474 |

**Fig. 2.**
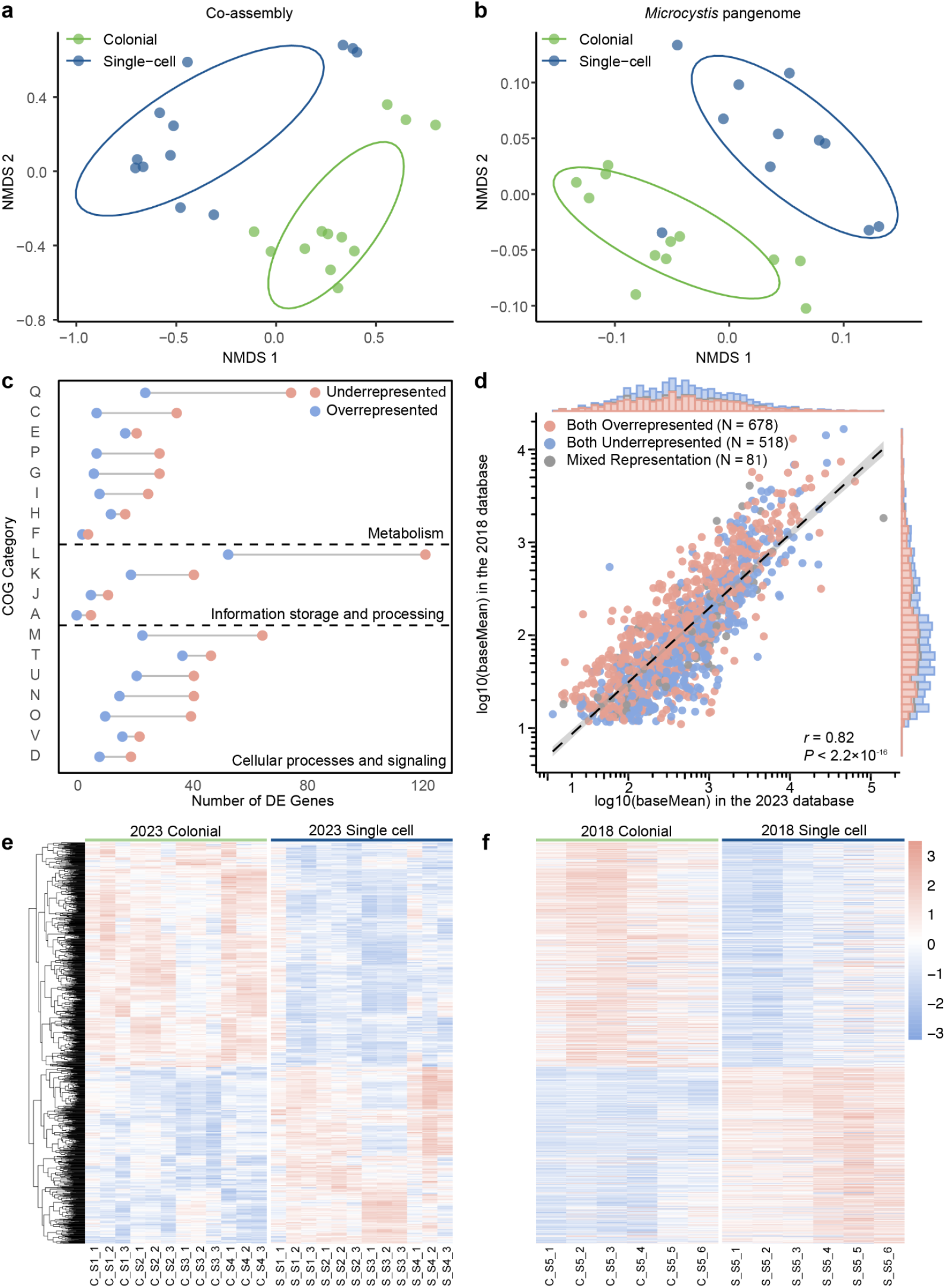
Functional distribution based on the co-assembly and *Microcystis* pangenome. **a, b**, Nonmetric multidimensional scaling (NMDS) plot based on Bray–Curtis dissimilarity of transcriptomic data mapped to the co-assembly (**a**) and the *Microcystis* pangenome (**b**), using transcript per million (TPM) normalized data. Ellipses cover 68% of the data for each form. (**c**), Distribution of 2166 differentially expressed (DE) *Microcystis* genes across COG functional categories. (**d**), Pearson correlation analysis of log10-transformed baseMean values for 1277 shared significant genes in the 2018 and 2023 datasets. (**e**), (**f**), Heatmap of 1277 shared significant genes across colonial and single-cell samples from the 2023 (**e**) and 2018 (**f**) dataset. Gene expression values were normalized to z-scores based on TPM values, and hierarchical clustering was performed on genes in the 2023 dataset, with the 2018 dataset using the same order.

To further investigate how community morphology contributed to the dissimilarity in *Microcystis* expression, DE gene analyses were performed based on the *Microcystis* pangenome and 2023 dataset. In total, 2,166 genes belonging to *Microcystis* were differentially expressed (|Log_2_ FC| ≥ 1, *p*_*adj*_ < 0.05), with 1,547 of these genes increased in relative expression in colonial samples and 619 decreased. Across each COG category, most genes exhibited increased expression in colonial samples (Fig. 2c). Genes categorized in COG category L (Replication, recombination and repair, n = 175) were the most highly represented category based in the DE dataset, with 69.71% of them overrepresented in colonial samples. Of those genes, the majority belonged to transposase-encoding genes, with 64 identified, of which 52 (81.25%) showed decreased expression in colonial samples (Supplementary Figure 3). Other genes associated with mobile genetic elements, such as genes encoding endonuclease and reverse transcriptase, also showed increased relative expression in colonies. Likewise, relative expression of genes within COG category Q (Secondary metabolites biosynthesis, transport and catabolism) increased in colonies (Supplementary Figure 3). Other genes, including those encoding PEP-CTERM protein, gas vesicles and calcium-binding protein exhibited increased expression in colonial samples (Supplementary Figure 3).

To confirm our 2023 observations, the 2018 dataset of twelve libraries was also examined to determine whether there DE genes from *Microcystis*. We identified 1,277 shared genes (adjusted *p* value of < 0.05) present in both datasets (summary of these genes are provided in Supplementary Data 1), and 93.7% of the genes showed consistent regulated results in two datasets. Correlation analysis (Fig. 2d) based on the average expression of shared genes in the two datasets revealed a strong positive correlation (Pearson correlation, *r* = 0.82, *p* < 0.001). We generated a gene clustering heatmap (Fig. 2e) using the 2023 dataset. Using the same gene order, we constructed a corresponding heatmap with the 2018 dataset (Fig. 2f). The results revealed consistent trends across both datasets, demonstrating that the significantly expressed genes exhibited the same regulatory patterns in both datasets. This indicated that the functional expression differences in *Microcystis* that are associated with differences in morphology were reproducible.

### Differential expression of genes from Microcystis-infecting phage

Based on *Microcystis* dominating both colonial and single-cell samples, as well as the functional differences driven by these morphological distinctions, we examined the expression of *Microcystis* phage genes to explore infection dynamics. Ma-LMM01, with its fully sequenced genome^35^, has served as an important model for studying *Microcystis*-phage interactions. Normalized expression of the Ma-LMM01 markers of lytic infection (tail sheath, *gp091*)^18^, and lysogenic infection (transposase, *gp135*, and site-specific recombinase, *gp136*)^18^ observed in colonial and single-cell samples are shown in Fig. 3a. Of the twenty-four samples from the 2023 dataset, significant differences in infection strategies were observed. Colonial samples showed lower expression of *gp091* (*p* < 0.001, paired *t*-test) and higher expression of *gp135* and *gp136* than single-cell samples (*p* < 0.001, paired *t*-test). In colonial samples, *gp135* and *gp136* exhibited significantly higher expression levels compared to *gp091* (Fig. 3c), with *gp135* showing a 1.64-fold increase (*p* = 0.012, paired *t*-test) and *gp136* displaying a 2.65-fold increase (*p* < 0.001, paired *t*-test), implying that lysogenic infection was dominant. The single-cell samples showed *gp091* expression to be 3.84 times (*p* = 0.0024, paired *t*-test) greater than *gp135* and 3.37 times (*p* = 0.027, paired *t*-test) greater than *gp136* (Fig. 3c), implying those free single *Microcystis* cells were experiencing more lytic infection. The expression of these maker genes was also examined in the 2018 dataset (Fig. 3b, d), revealing a consistent trend but with even greater difference. These expression pattern differences were consistent across different collection years, lake locations, and times of day.

**Fig. 3.**
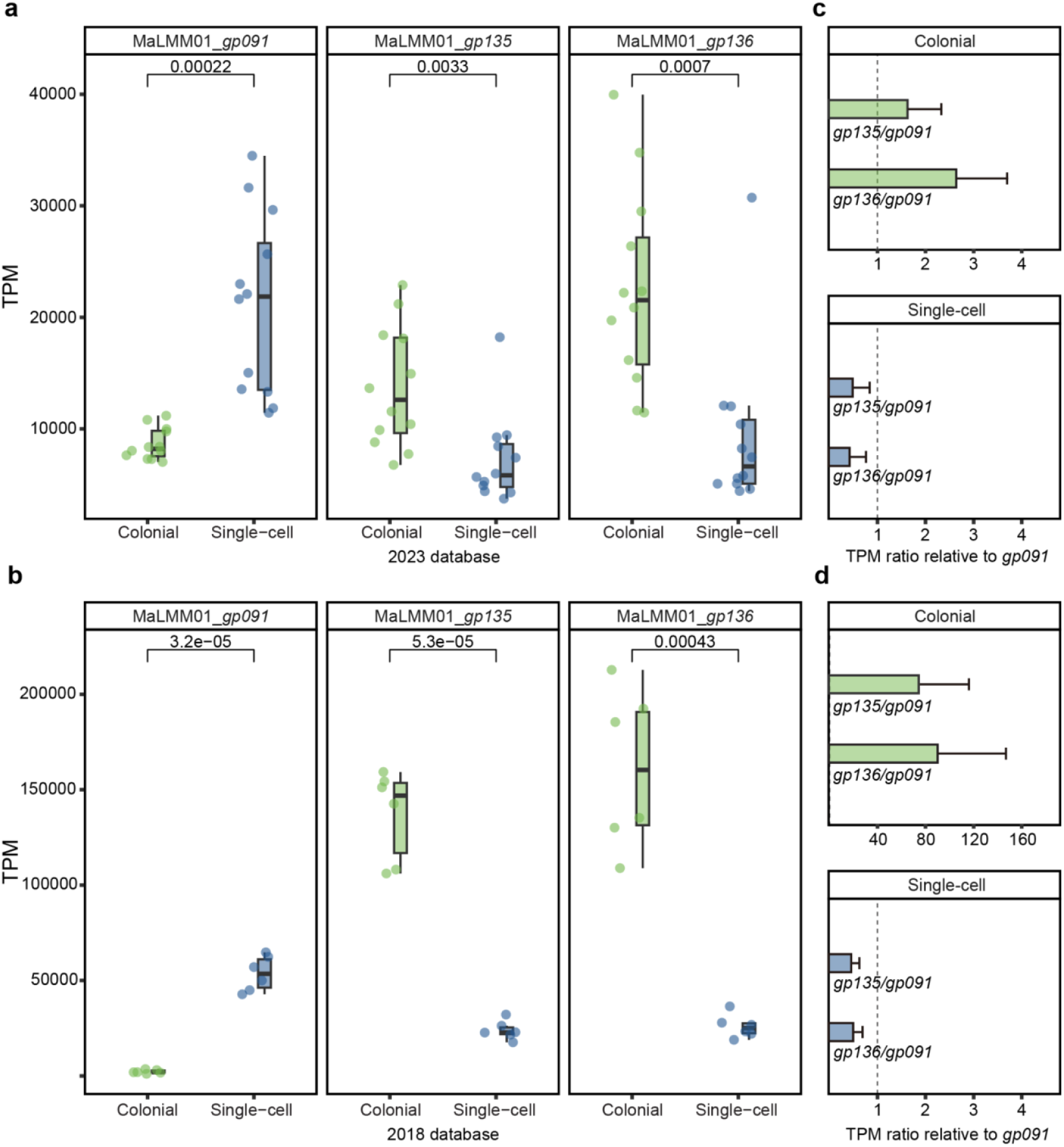
Relative expressions of lysogeny- and lysis-associated genes. **a, b**, TPM values of lysogeny- and lysis-associated genes in colonial and single-cell samples from the 2023 (**a**) and 2018 (**b**) dataset. *gp091* represents a marker for lytic infection, while *gp135* and *gp136* represent markers for lysogenic infection. The horizontal bars within the boxes represent medians, while the tops and bottoms of the boxes indicate the 75th and 25th percentiles, respectively. The upper and lower whiskers extend to the furthest data points within 1.5× the interquartile range from the edges of the box. **c, d**, TPM ratio of lysogeny-associated genes (*gp135* and *gp136*) relative to the lysis-associated gene (*gp091*) in colonial and single-cell samples from the 2023 (**a**) and 2018 (**b**) dataset. Ratios greater than 1 indicate lysogeny-dominant activity, while ratios less than 1 indicate lysis-dominant activity. Error bars represent the standard error of the mean (SEM) for each ratio. The dashed vertical line at 1 represents equal dominance between lysogeny and lysis.

**Fig. 4.**
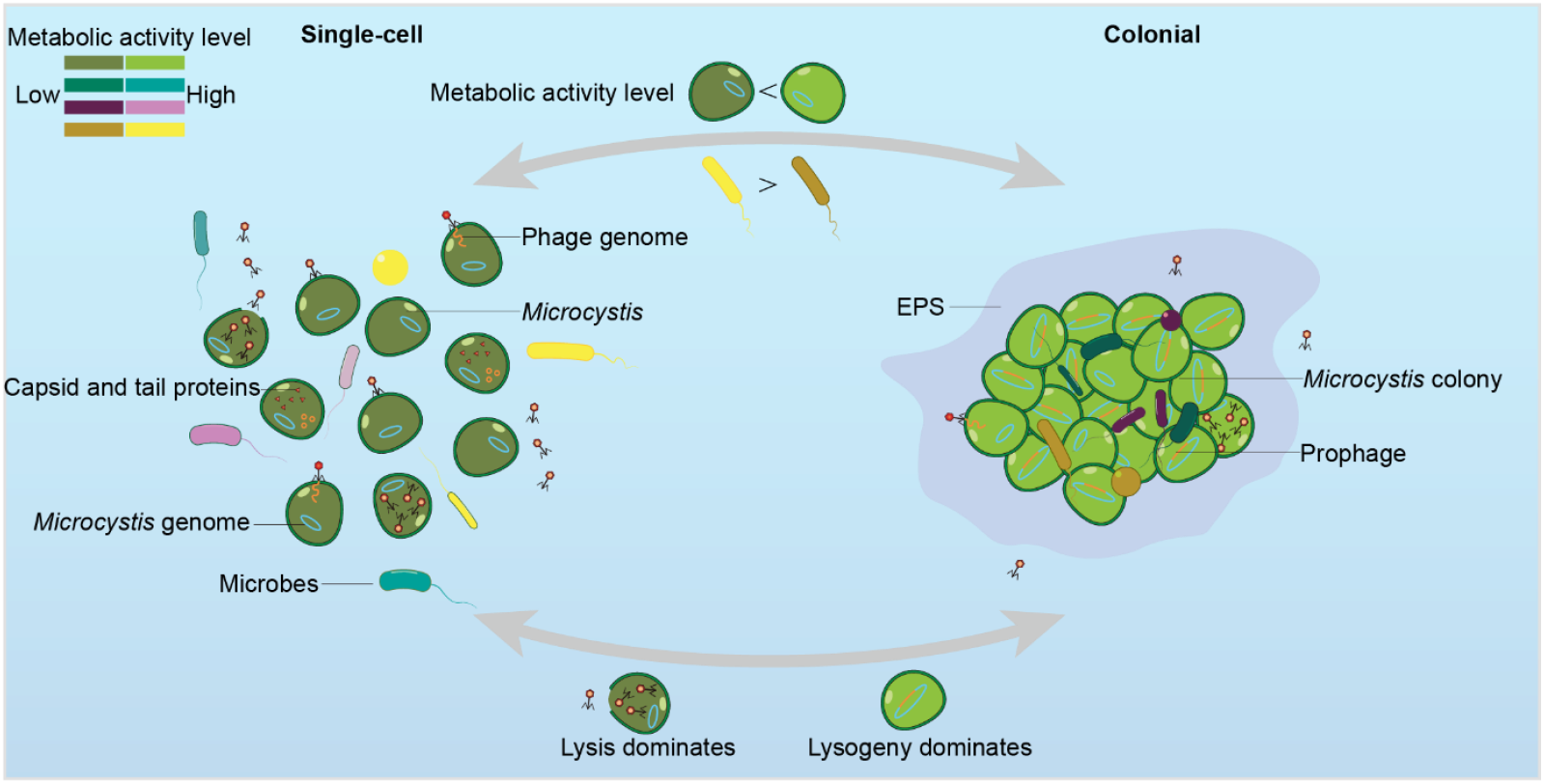
Metabolic activity and phage-host interactions in single-cell and colonial *Microcystis*. The left panel shows single-cell *Microcystis*, characterized by higher activity in associated microbes and lytic phage dominance. The right panel represents colonial *Microcystis*, where metabolic activity is concentrated in *Microcystis* and lysogeny dominates. Arrows indicate transitions between single-cell and colonial forms, driven by environmental factors and phage infection dynamics. Metabolic activity levels are represented using color brightness (Bright: high activity; Dark: low activity).

## Discussion

This study focused on the dominant genus of cyanobacterial blooms in Lake Taihu, *Microcystis*, and explored its community dynamics, interactions with associated bacteria, and the infection strategies of *Microcystis* phages. Using metatranscriptomic data collected in 2023, we examined differences in species abundance and functional activity across colonial and single-cell forms of *Microcystis* and its associated microbes, as well as the contrasting infection strategies of *Microcystis* phages in these two forms. The findings were validated using a similar 2018 metatranscriptomic dataset.

At each location, samples of both colonial and single-cell forms were collected in 2023 under identical environmental conditions, including temperature and nitrogen and phosphorus concentrations. This consistency ensures that observed differences in the taxonomic structure and functional composition of aquatic microbial communities were primarily attributable to the formation of colonies. The differences were likely driven by the dominant role of *Microcystis* within the colonies and its interactions with associated microbes.

Colonial *Microcystis* is sometimes considered to be more competitive than the single-cell form due to enhanced resilience to environmental stresses provided by its structural advantages, such as the extracellular polymeric substance (EPS) matrix that offers physical protection and facilitates nutrient acquisition^6^. Colony formation is also thought to shield individual cells from environmental pressures^6^. However, overrepresentation of many other transcripts in colonial *Microcystis* suggests that significant metabolic investment is required to maintain a colony. This duality implies a trade-off: while colonies may offer greater defense and stability, they appear to demand higher metabolic effort, which ultimately shapes the competitive fitness of colonial *Microcystis*.

Interestingly, *Microcystis* genes exhibit increased expression in colonies, while the transcripts of associated microbes were reduced in representation. This suggests that microbiome members may need to be maintain more active metabolic pathways in single-cell environments. We hypothesize that during the transition from single-cell to colonial *Microcystis*, microbial interactions undergo significant changes. In the initial stages of colony formation, microbes may collaborate but rely on their individual metabolic pathways. Once the colony is established, the system aligns with the “Black Queen Hypothesis”^36^: *Microcystis* assumes a central role, taking on the majority of metabolic and defensive burdens, while associated microbes downregulate gene expression, benefiting from a stable and protective environment where metabolic burdens are assumed by the cyanobacterium. To this end, while some researchers have hypothesized that blooms form and persist due to the activity of associated microbial community members^37^, it appears that the opposite may in part be true: blooms and particularly colonies provided a haven for many heterotrophic bacteria, perhaps through both the increased physical protection from the colony as well as the surplus of metabolic products *Microcystis* appears to produce. Given the caveats of metatranscriptomic work (the often quoted “*transcription is not translation*”, as well as the numerical differences in *Microcystis* and microbiome members, this emergent hypothesis necessitates future study.

Perhaps the most unique observation from this data set is the striking difference between potential lysogens and lytically-infected *Microcystis* cells in the colonial *vs* single cell samples. Studies on lysogen formation in *Microcystis* are in their infancy. But, the possibility of lysogen formation by Ma-LMM01-like phages was noted with the initial genomic sequencing of the virus^35^, and field studies in *Taihu* demonstrated strong seasonal patterns in expression of genes to which lysogenic function is ascribed^18^. However, the factors promoting or constraining these infection outcomes (and indeed even which partner makes that decision) are unclear. In the present case, lysogen formation appears to be consistent with the hypothesis that this relationship forms/is selected for during life-at-high-density scenarios^38^: many lysogens demonstrate immunity to superinfection by similar viruses. During large scale *Microcystis* blooms, both the host and its virus can reach densities greater than 100,000 per ml^39^. At these densities viruses should contact potential hosts on a daily basis^19^, effectively collapsing the bloom as is seen in other high density algal blooms^40^. Given that colonies are a localized high density scenario, protection from superinfection would seem to be a necessary priority for members of the colony forming community. Going forward, it will be interesting to determine how colonies can mimic climax bloom communities as opposed to early season populations, which likely mimic single cells^41^.

## Conclusion

Our study illuminates how colony formation restructures *Microcystis*’ metabolic activity and alters its interactions with both associated bacteria and phages. Colonies appear to harbor a specialized microenvironment where *Microcystis* invests in sustaining a dense population, maintains putative fitness benefits such as enhanced stress tolerance, and supports an elevated propensity for lysogenic phage infection. By contrast, single-cell *Microcystis* undergoes more frequent lytic attacks, likely due to reduced cell density and diminished collective immunity. These findings highlight the dual ecological role of colony formation as both a protective refuge and a driver of complex host–virus dynamics.

Future controlled experiments—whereby environmental parameters, *Microcystis* colony density, and phage populations are systematically manipulated—will be essential to pinpoint the mechanistic underpinnings of lysogenic switching and to clarify the interplay between host competition, viral infection modes, and bloom persistence. Given the potentially key role of viruses in releasing toxins from the particulate to dissolved fraction in aquatic environments^42,43^, understanding these processes are key to understanding system ecology and protecting our water resources. Cyanobacterial blooms and their associated toxins pose significant risks to aquatic ecosystems, drinking water supplies, and public health. By shedding light on these interactions, we can improve predictive modeling for bloom development and advance more effective strategies to mitigate their effects. Ultimately this would improve the availability of safe and sustainable water resources for ecosystems and human use alike.

## Acknowledgements

We are grateful to Zunqing Du for her help with 2023 sample collection as well as the scientists and staff of the *Taihu Laboratory for Lake Ecosystem Research* for assistance with the 2018 collections. A portion of the computation for this work was performed on the University of Tennessee *Infrastructure for Scientific Applications and Advanced Computing (ISAAC)* computational resources.

