## Supplementary Tables and Figures for "Lysogen formation governs colonies while lytic infection is more prevalent in single cells of the bloom-forming cyanobacterium, *Microcystis*"

METHODS

#### Collection of 2018 samples

Surface water samples were collected from a dock that stretches approximately 100 yards into Meilang Bay near the TLLER field station on *Taihu* on 08/18/2018 at 13:00 and 08/19/2018 at 7:00. To enrich for colonies, water was concentrated on a 28 µm mesh while the < 28 µm filtrate was captured as the representative single-cell community. Duplicate samples were collected at each time point, and both fractions were filtered through 0.2 µm Sterivex, preserved with RNA*later* and stored at -80° C until processing.

#### RNA extraction and sequencing of 2018 samples

Total RNA was extracted using a previously described phenol–chloroform method followed by ethanol precipitation[1]. Genomic DNA was removed using the Turbo DNA-free kit (Ambion, Austin, TX, USA). RNA integrity and concentration were assessed with the Qubit RNA HS Assay Kit (Invitrogen, Waltham, MA, USA). Samples exhibiting residual DNA, as indicated by Qubit quantification, underwent an additional DNase treatment. Purified RNA was subsequently subjected to library preparation, rRNA depletion, and sequencing on the Illumina NextSeq platform using the Epidemiology Ribo-Zero Gold rRNA Removal Kit.

| Sampling site | Date | Time | Longitude | Latitude | Colonial samples | Single-cell samples |
|---|---|---|---|---|---|---|
| S1 | 8/26/2023 | 10:02 | 120°01’21’’E | 31°24’48’’N | C_S1_1<br>C_S1_2<br>C_S1_3 | S_S1_1<br>S_S1_2<br>S_S1_3 |
| S2 | 8/26/2023 | 10:13 | 120°00’10’’E | 31°23’28’’N | C_S2_1<br>C_S2_2<br>C_S2_3 | S_S2_1<br>S_S2_2<br>S_S2_3 |
| S3 | 8/26/2023 | 10:27 | 119°58’30’’E | 31°20’55’’N | C_S3_1<br>C_S3_2<br>C_S3_3 | S_S3_1<br>S_S3_2<br>S_S3_3 |
| S4 | 8/26/2023 | 11:15 | 120°02’43’’E | 31°23’51’’N | C_S4_1<br>C_S4_2<br>C_S4_3 | S_S4_1<br>S_S4_2<br>S_S4_3 |

Supplementary Table 1: Information of each sampling site. The first letter of each sample name indicates the morphology of the sample, with C representing colony and S representing single cell. The second segment represents the sampling site (e.g., S2), and the last number indicates the replicate number.

| Organism Scientific Name | Organism Qualifier | Taxonomy id | Assembly Accession | Source | Size | Gene Count |
|---|---|---|---|---|---|---|
| Microcystis aeruginosa NIES-843 | strain: NIES-843 | 449447 | GCF_000010625.1 | RefSeq | 5842795 | 5792 |
| Microcystis aeruginosa NIES-2549 | strain: NIES-2549 | 1641812 | GCF_000981785.2 | RefSeq | 4301200 | 4087 |
| Microcystis panniformis FACHB-1757 | strain: FACHB-1757 | 1638788 | GCF_001264245.1 | RefSeq | 5686839 | 5640 |
| Microcystis aeruginosa NIES-2481 | strain: NIES-2481 | 1698524 | GCF_001704955.2 | RefSeq | 4440545 | 4242 |
| Microcystis aeruginosa PCC 7806SL | strain: PCC 7806SL | 1903187 | GCF_002095975.1 | RefSeq | 5139339 | 4936 |
| Microcystis sp. MC19 | strain: MC19 | 1967666 | GCF_003019735.1 | RefSeq | 5020243 | 4934 |
| Microcystis viridis NIES-102 | strain: NIES-102 | 213615 | GCF_003945305.1 | RefSeq | 5874197 | 5786 |
| Microcystis aeruginosa FD4 | strain: FD4 | 2686288 | GCF_009792235.1 | RefSeq | 5493112 | 5530 |
| Microcystis aeruginosa NIES-298 | strain: NIES-298 | 449468 | GCF_010196425.1 | RefSeq | 5015081 | 4823 |
| Microcystis aeruginosa | strain: NIES-88 | 1126 | GCF_019704275.1 | RefSeq | 5501105 | 5588 |
| Microcystis aeruginosa FACHB-905 = DIANCHI905 | strain: DIANCHI905 | 267865 | GCF_021172085.1 | RefSeq | 5103104 | 4909 |
| Microcystis aeruginosa str. Chao 1910 | strain: Chao 1910 | 2945101 | GCF_026222535.1 | RefSeq | 5669822 | 5720 |
| Microcystis aeruginosa PCC 7806 | strain: PCC 7806 | 267872 | GCF_030553035.1 | RefSeq | 5103923 | 4908 |
| Microcystis aeruginosa NRERC-214 | strain: NRERC-214 | 2528657 | GCF_031754835.1 | RefSeq | 4981678 | 4671 |
| Microcystis aeruginosa 1339 | strain: 1339 | 3113711 | GCF_038396575.1 | RefSeq | 4749293 | 4559 |
| Microcystis aeruginosa PCC 7806 | strain: PCC 7806 | 267872 | GCF_041506625.1 | RefSeq | 5096229 | 4894 |

Supplementary Table 2: Summary of 16 complete *Microcystis* genomes used for the construction of the *Microcystis* pangenome.

| Sample | Raw reads count | Total bases of raw reads | Trimmed reads count | Trimmed reads percentage (%) | mRNA reads count | mRNA reads percentage (%) | Reads mapped to the *Microcystis* pangenome count | *Microcystis* pangenome -mapped reads percentage (%) | Reads mapped to the co-assembly count | Co-assembly-mapped reads percentage (%) |
|---|---|---|---|---|---|---|---|---|---|---|
| C_S1_1 | 1.55E+08 | 2.33E+10 | 1.52E+08 | 98.17% | 1.52E+08 | 99.93% | 5.53E+07 | 36.49% | 1.34E+08 | 88.29% |
| C_S1_2 | 1.58E+08 | 2.39E+10 | 1.54E+08 | 97.50% | 1.54E+08 | 99.92% | 6.85E+07 | 44.43% | 1.4E+08 | 90.65% |
| C_S1_3 | 9.74E+07 | 1.47E+10 | 9.53E+07 | 97.86% | 9.52E+07 | 99.95% | 3.32E+07 | 34.9% | 8.39E+07 | 88.11% |
| C_S2_1 | 1.4E+08 | 2.12E+10 | 1.37E+08 | 97.95% | 1.37E+08 | 99.75% | 5.13E+07 | 37.41% | 1.23E+08 | 90.03% |
| C_S2_2 | 1.18E+08 | 1.78E+10 | 1.15E+08 | 97.57% | 1.15E+08 | 99.71% | 4.37E+07 | 38.1% | 1.03E+08 | 89.79% |
| C_S2_3 | 8.99E+07 | 1.36E+10 | 8.78E+07 | 97.69% | 8.78E+07 | 99.96% | 2.79E+07 | 31.79% | 7.83E+07 | 89.16% |
| C_S3_1 | 1.4E+08 | 2.11E+10 | 1.32E+08 | 94.90% | 1.31E+08 | 99.05% | 5.95E+07 | 45.38% | 1.21E+08 | 92.48% |
| C_S3_2 | 1.67E+08 | 2.52E+10 | 1.63E+08 | 97.77% | 1.63E+08 | 99.84% | 7.31E+07 | 44.89% | 1.5E+08 | 91.97% |
| C_S3_3 | 9.68E+07 | 1.46E+10 | 9.58E+07 | 98.98% | 9.43E+07 | 98.35% | 4.3E+07 | 45.6% | 8.61E+07 | 91.31% |
| C_S4_1 | 1.35E+08 | 2.04E+10 | 1.28E+08 | 94.87% | 1.28E+08 | 99.94% | 5.61E+07 | 43.8% | 1.17E+08 | 91.09% |
| C_S4_2 | 1.08E+08 | 1.62E+10 | 1.06E+08 | 98.37% | 1.06E+08 | 99.90% | 4.17E+07 | 39.45% | 9.66E+07 | 91.36% |
| C_S4_3 | 9.6E+07 | 1.45E+10 | 9.36E+07 | 97.44% | 9.35E+07 | 99.95% | 3.42E+07 | 36.54% | 8.3E+07 | 88.75% |
| S_S1_1 | 1.15E+08 | 1.74E+10 | 1.12E+08 | 97.78% | 1.12E+08 | 99.94% | 3.59E+07 | 31.95% | 9.83E+07 | 87.52% |
| S_S1_2 | 1.14E+08 | 1.73E+10 | 1.11E+08 | 97.19% | 1.11E+08 | 99.92% | 2.21E+07 | 19.93% | 8.98E+07 | 80.82% |
| S_S1_3 | 1.14E+08 | 1.72E+10 | 1.1E+08 | 96.68% | 1.1E+08 | 99.89% | 2.96E+07 | 26.86% | 9.32E+07 | 84.52% |
| S_S2_1 | 1.13E+08 | 1.7E+10 | 1.1E+08 | 97.07% | 1.09E+08 | 99.87% | 1.62E+07 | 14.84% | 8.46E+07 | 77.29% |
| S_S2_2 | 1.08E+08 | 1.63E+10 | 1.06E+08 | 98.21% | 1.06E+08 | 99.93% | 1.27E+07 | 11.99% | 8.49E+07 | 80.36% |
| S_S2_3 | 1.02E+08 | 1.54E+10 | 9.99E+07 | 98.08% | 9.97E+07 | 99.84% | 1.72E+07 | 17.25% | 8.53E+07 | 85.53% |
| S_S3_1 | 1.37E+08 | 2.07E+10 | 1.34E+08 | 97.43% | 1.34E+08 | 99.95% | 2.7E+07 | 20.19% | 1.18E+08 | 88.27% |
| S_S3_2 | 1.46E+08 | 2.21E+10 | 1.43E+08 | 97.79% | 1.43E+08 | 99.93% | 3.35E+07 | 23.41% | 1.28E+08 | 89.4% |
| S_S3_3 | 1.36E+08 | 2.06E+10 | 1.33E+08 | 97.52% | 1.33E+08 | 99.93% | 3.5E+07 | 26.36% | 1.19E+08 | 89.36% |
| S_S4_1 | 1.26E+08 | 1.9E+10 | 1.22E+08 | 96.46% | 1.21E+08 | 99.87% | 1.48E+07 | 12.19% | 9.7E+07 | 79.93% |
| S_S4_2 | 1.38E+08 | 2.08E+10 | 1.33E+08 | 96.26% | 1.32E+08 | 99.83% | 1.73E+07 | 13.03% | 1.08E+08 | 81.65% |
| S_S4_3 | 1.02E+08 | 1.54E+10 | 9.95E+07 | 97.68% | 9.94E+07 | 99.91% | 1.07E+07 | 10.73% | 8.02E+07 | 80.61% |

Supplementary Table 3: Summary of sequence information for Lake Taihu metatranscriptomic libraries collected on August 26, 2023. The first letter of each sample name indicates the morphology of the sample, with C representing colony and S representing single cell. The second segment represents the sampling site (e.g., S2), and the last number indicates the replicate number.

#### Supplementary Figure 1

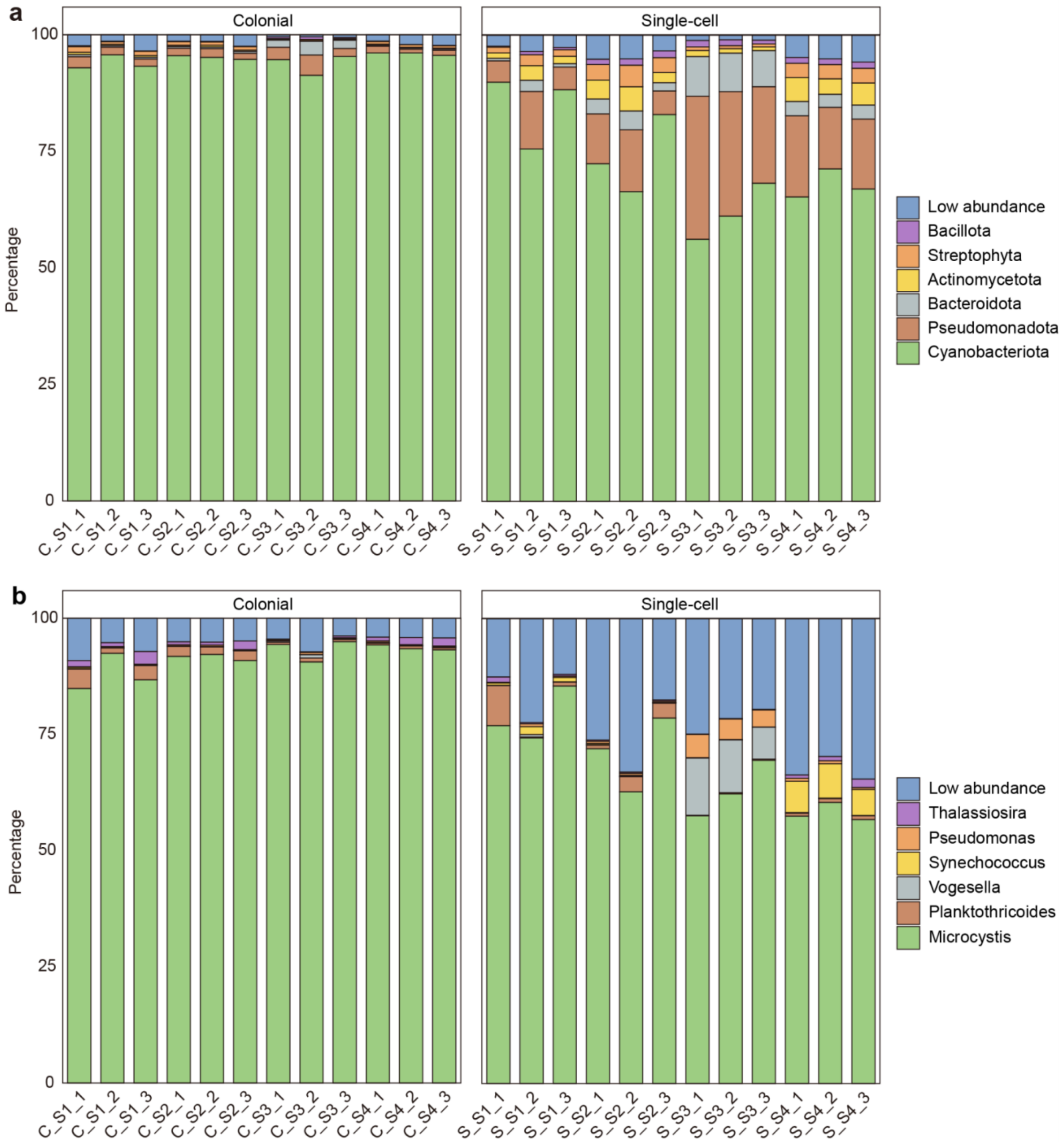


Supplementary Figure 1: Phylum- (a) and genus-level (b) distribution of colonial and single-cell samples in the 2023 database. Each bar represents a single sample, with the relative abundance of taxa shown as a percentage. The number of samples is as follows: colonial (n = 12) and single-cell (n = 12).

### Supplementary Figure 2

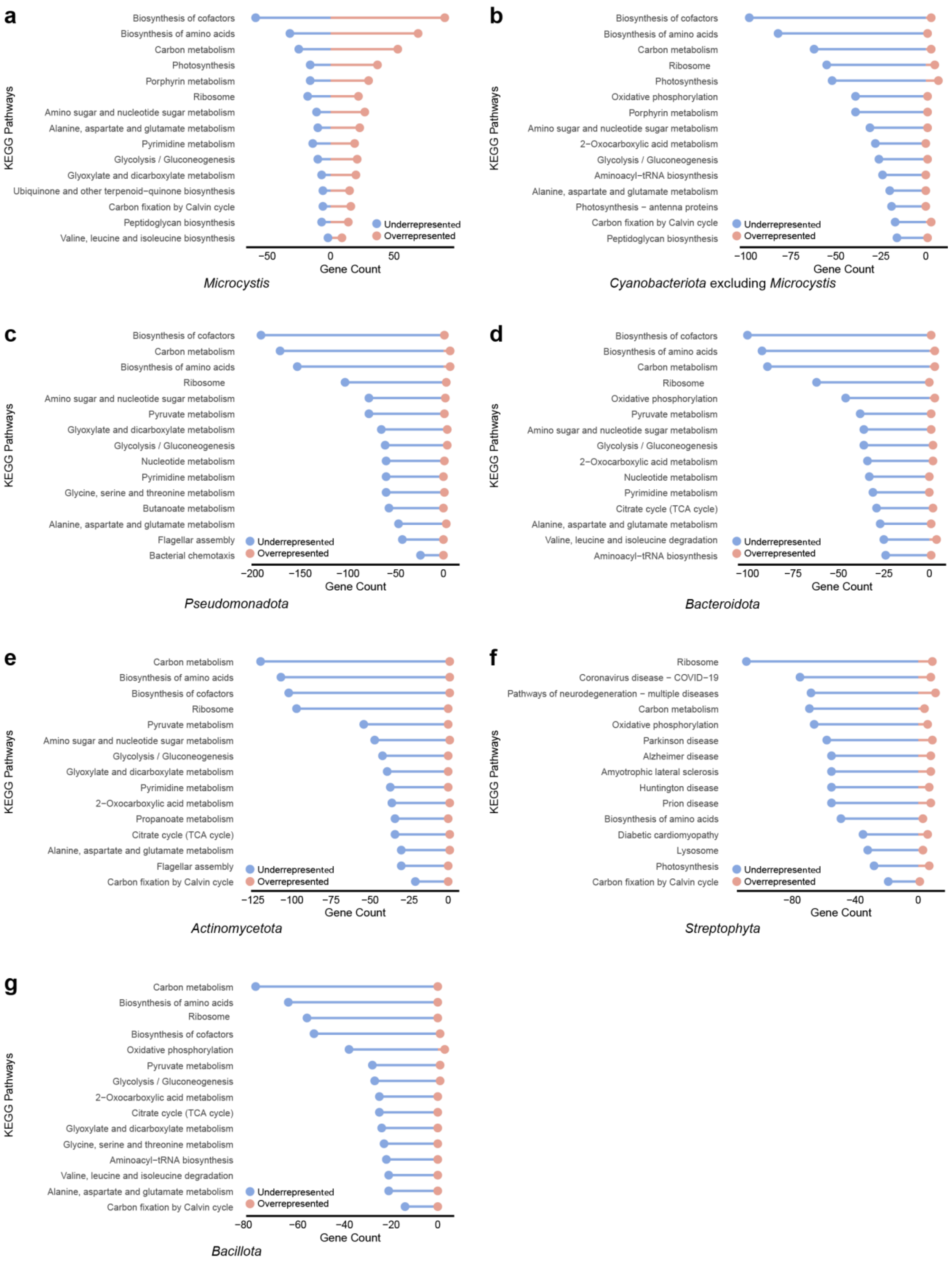

**Supplementary Figure 2:** Top 15 KEGG pathways enriched in the top six phyla based on differential gene expression analysis. The phyla include **a** *Microcystis*, **b** *Cyanobacteriota* (excluding *Microcystis*), **c** *Pseudomonadota*, **d** *Bacteroidota*, **e** *Actinomycetota*, and **g** *Bacillota*. Each panel shows the number of overrepresented (red) and underrepresented (blue) genes in key metabolic and functional pathways. KEGG pathways are ranked by gene count. *Cyanobacteriota* was divided into *Microcystis* and non-*Microcystis Cyanobacteriota*.

#### Supplementary Figure 3

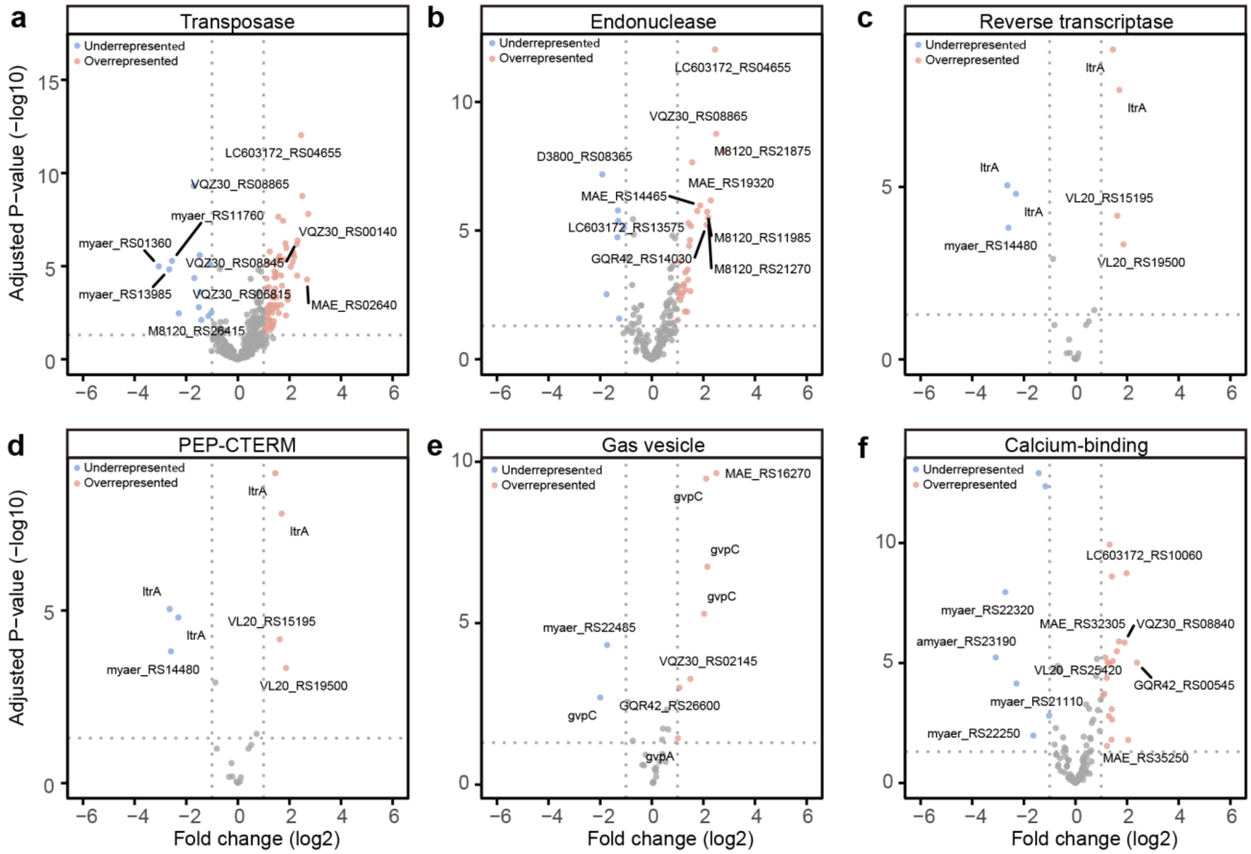


**Supplementary Figure 3:** Volcano plots depicting the differential expression and significance of genes across colonial and single-cell samples. **a**, **b**, and **c** highlight key genes categorized under COG category L, specifically transposase (**a**), endonuclease (**b**), and reverse transcriptase (**c**). **d**, **e**, and **f** present genes without COG classification, showcasing PEP-CTERM (**d**), gas vesicle proteins (**e**), and calcium-binding proteins (**f**). Red and blue points denote overrepresented and underrepresented genes, respectively, while gray points represent non-significant genes (adjusted p-value > 0.05). Annotated points indicate specific genes of interest.

1. Zepernick, B. N. *et al.* Declines in ice cover are accompanied by light limitation responses and community change in freshwater diatoms. *ISME J.* **18**, wrad015 (2024).
